# Hydrolysis-dependent severing tunes internal monomeric heterogeneity to shape actin length distributions

**DOI:** 10.1101/2025.05.29.656816

**Authors:** Soumyadipta Ray, Binayak Banerjee, Lishibanya Mohapatra, Dipjyoti Das

**Author notes:** contributed equally (Soumyadipta Ray, Binayak Banerjee).

## Abstract

The coordination of actin-binding proteins (ABPs) is essential for forming a wide range of actin structures that drive vital cellular processes. Many of the ABPs, like the severing protein cofilins, exhibit affinities dependent on the nucleotide state of actin monomers. Experiments have shown that cofilins interact with actin filaments cooperatively in a concentration-dependent manner, where it severs less at both low and high concentrations. However, how this nucleotide state-dependent severing influences actin filament distributions is poorly understood. Here, we propose a computational model of actin filament dynamics that combines stochastic polymerization, phosphate release, hydrolysis-dependent cooperative binding of cofilins, and severing of actin monomers. This model simultaneously reproduces key experimentally observed features: The non-monotonic change of the filament length and the monotonic decay in actin cap size with the increasing cofilin concentration, as well as the linear increase of the cap size with the actin concentration. We elucidate how both these emerge from the tuning of the heterogeneity in the arrangement of cofilin decorated and undecorated monomers within a filament. We also predict that, with varying cofilin concentrations, steady-state length distributions can change from symmetric bell-shaped to long-tailed skewe d shapes. We show that, in some regimes, our model simplifies to a coarse-grained model enabling us to mathematically predict that the distribution of undecorated monomers is exponential with a decay exponent that depends on cofilin and actin concentration. This study highlights the importance of non equilibrium processes that can modulate structural heterogeneity within a filament to shape actin length variation.

## INTRODUCTION

Actin filaments are a crucial component of the eukaryotic cytoskeleton, supporting essential cellular processes such as cell motility, division, vesicle transport, molecular motor movement, and the maintenance of cell shape (1). Actin-based structures are highly variable, ranging from actin patches to cables (2). Their dynamics differ significantly — actin networks at endocytic sites and lamellipodia are short-lived, lasting only a few seconds, whereas actin bundles in structures like filopodia, stereocilia, and microvilli can remain stable for minutes to days (3). This structural variability highlights significant variation in the size and dynamics of actin assemblies.

Experimental evidence suggests that this variability in actin dynamics and rapid turnover within cells is driven by the spatiotemporal coordination of multiple actin-binding proteins (ABPs) (4). Although key ABPs that regulate actin assembly and disassembly are known (5), how the regulatory proteins ultimately affect the emerging lengths of actin filaments and their size variation remains a fundamental challenge in cell biology. Among the ABPs, the actin depolymerizing factor (ADF), cofilins, can sever actin filaments in a concentration-dependent manner, promoting random and rapid disassembly of filaments (6–9). These severing proteins are expressed in all eukaryotic cells (10) and are involved in a range of cellular functions, including motility (11, 12) and hence the focus of our study.

Many in vitro experiments have explored the effect of severing proteins on actin filaments. For instance, Pavlov et al. (7) studied the ADF/cofilin interactions with actin filaments and found that cofilin-dependent severing of actin filaments first increased and then gradually decreased with increasing cofilin concentration (see Fig 1A). They also reported skewed long-tailed length distributions at several cofilin concentrations, suggesting that huge length fluctuations may emerge due to severing. Another experiment (8) also reported a concentration dependence in cofilin’s severing activity. The authors found that at very low concentrations, cofilins do not bind to actin filaments. At an optimal intermediate concentration, cofilins bind and sever actin filaments. In contrast, at higher concentrations, cofilins bind cooperatively to actin filaments and promote ATP hydrolysis but hardly sever them. These experiments suggest that the actin filament length changes non-monotonically with the cofilin concentration.

**Fig 1:**
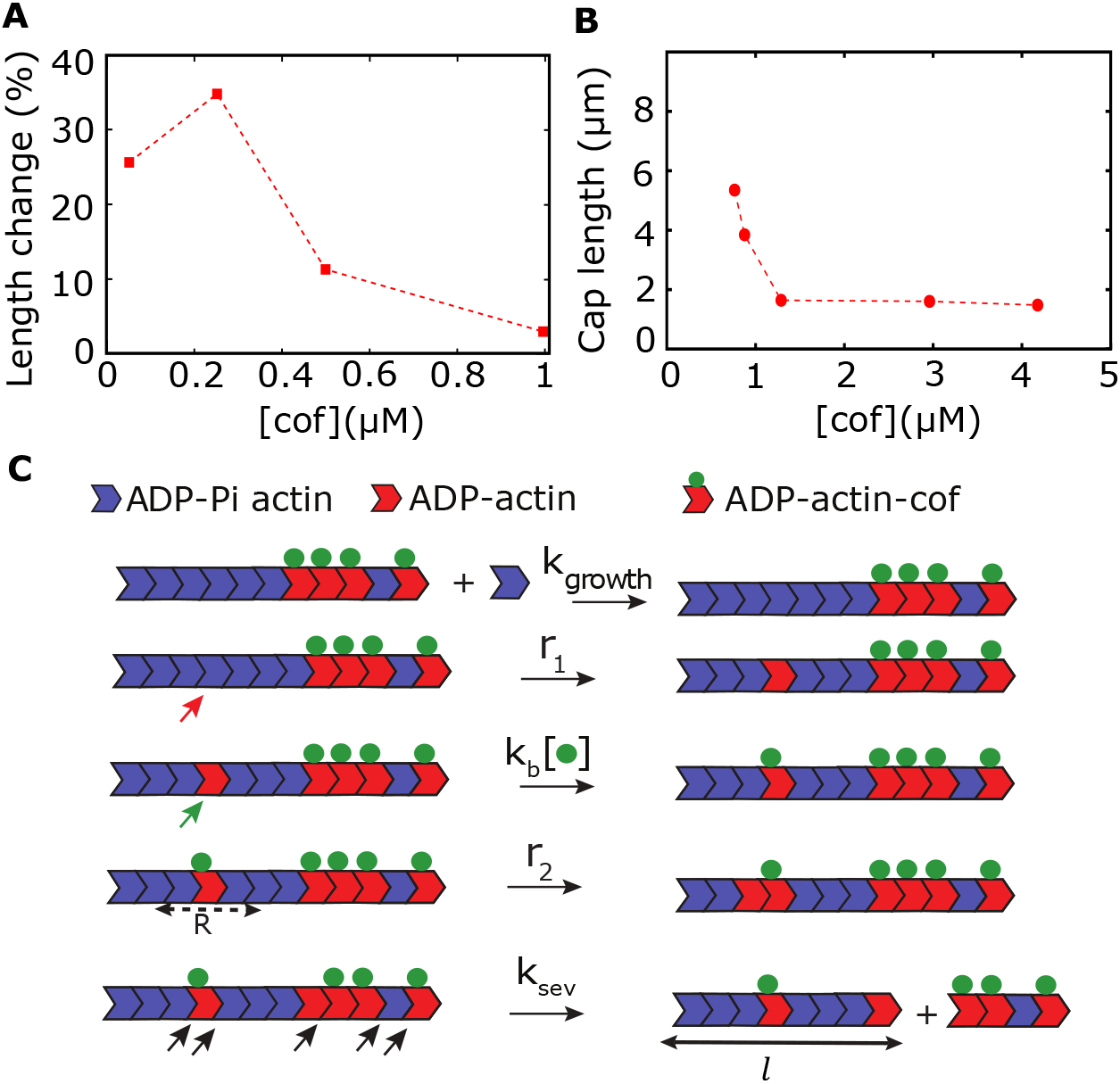
A three-state model of hydrolysis-dependent severing. **(A-B)** Published experimental data report (A) a non-monotonic change in the mean length of actin filaments (7) and (B) a monotonic decrease in the ATP cap length of actin filaments with cofilin concentration, denoted by [*cof*] (9). **(C)** Our proposed model of the hydrolysis-dependent severing of actin filament consists of three states of monomer: ADP-Pi actin (blue), ADP actin (red), and cofilin-bound ADP actin, which undergo five processes: growth (with rate *k*_*growth*_), Pi release from actin monomers within a filament (with rate *r*_1_, denoted by the red slanting arrow), binding of the severing protein, cofilin, to ADP actin (*k*_*b*_[•], denoted by the green slanting arrow), faster Pi release (with rate *r*_2_ > *r*_1_) when the ADP-actin monomer is within a range *R* from a cofilin-bound ADP actin, and severing (with rate *k*_*sev*_, possible severing sites denoted by black slanting arrows). Note that, after severing, length is measured for the filament segment connected to the original barbed end (denoted by *l*). [•] denotes the cofilin concentration.

Some computational studies have previously explored the effect of severing on actin dynamics (13) and showed that cofilin severing can control actin filament lengths, producing a peaked length distribution. However, these studies neglected the hydrolysis-dependent severing activity. Generally, actin monomers are bound to ATP, and within the filaments, the ATP-bound monomers transform into ADP-bound monomers, releasing the phosphate (Pi) through hydrolysis. It was previously shown that distinct kinematic properties of ATP and ADP monomers produced by hydrolysis could lead to higher length fluctuations (14) and higher collective force generation by a bundle of growing filaments (15, 16) when compared to non-hydrolyzable filaments. Notably, cofilin binding to actin monomers can affect the hydrolysis, altering the Pi-release rate (6) and impacting the actin dynamics.

Depending on previous experimental observations (6), a computational model by Roland et al. (17) assumed that cofilins preferentially and cooperatively bind to ADP-monomers and accelerate the phosphate release of neighboring unhydrolyzed monomers. However, this model predicted bell-shaped length distributions unlike long-tailed distributions (i.e., high length fluctuations) found in experiments (7). This model also could not capture the non-monotonic behavior of actin lengths with the cofilin concentration. Further, this model assumed that the severing occurs between the adjacent cofilin-bound sites, which was later seen as not the case.

The experiments by Suarez et al.(9) further shed light on how severing depends on the nucleotide state of the monomers. Their experiments indeed indicate that bound cofilin accelerates Pi release from adjacent unoccupied monomers, effectively increasing the hydrolysis rate. Importantly, they found that severing occurs mainly at boundaries between bare and cofilin-decorated filament segments. By including this new observation, the authors modified the model by Roland et al.(17) to study the effect on actin length dynamics. In particular, they measured the length of actin caps (i.e., the continuous segment of unhydrolyzed actin monomers near the barbed end) and showed that the actin cap lengths monotonically and rapidly decrease with cofilin concentration (Fig 1B), in contrast with the non-monotonic change in the length of entire filaments (Fig 1A) (7). However, the authors did not explore the effect of hydrolysis-dependent severing on the filament length distribution and length variability.

In this paper, we introduce a coarse-grained model of actin filament dynamics to understand how the coupling between phosphate release and cofilin binding impacts the severing activity and resulting actin filament length distributions. Our model combines the random polymerization of actin monomers, hydrolysis-dependent cooperative binding of cofilins, cofilin-dependent enhancement of Pi release, and stochastic severing of actin segments. Our model simultaneously captures many experimentally observed features: The non-monotonic change of the filament length and the monotonic decay in actin cap size with the increasing cofilin concentration, as well as the linear increase of the cap size with the actin concentration. Moreover, using stochastic simulations, we predict the effect of severing on steady-state length distributions — the distributions change from bell-shaped to skewed long-tailed shapes with varying cofilin concentrations, passing through a minimum length fluctuation regime (quantified by the Fano factor). We further identify the limiting conditions where our model reduces to a simpler two-state coarse-grained model, allowing us to mathematically predict the average cap size and the distribution of cofilin-undecorated monomers along a filament at different actin and cofilin concentrations. We show that microscopic heterogeneity in the arrangement of decorated and undecorated monomers within a filament can give rise to macroscopic effects, such as variations in filament length and length distributions.

## MODEL

We develop a kinetic model to describe the dynamics of an actin filament undergoing the processes of polymerization, phosphate release, cooperative binding of cofilins, and severing of actin monomers. We consider that the actin filament polymerizes from the barbed end (Fig 1C). The actin monomers can bind to the barbed end with a rate of *k*_*growth*_ = *r*_0_ × [*actin*], where *r*_0_ is the bare rate of polymerization and [*actin*] denotes the actin concentration in the solution. Note that the actin monomers assembled into a filament are typically bound to ATP. These ATP monomers inside a filament irreversibly convert to ADP monomers via ATP hydrolysis, which is a two-step process: First, ATP monomers rapidly covert to ADP-Pi monomers via ATP cleavage, followed by a slower phosphate (Pi) release, producing ADP monomers (18). Nevertheless, since the slow step of Pi release is rate-limiting, we assume a simplified hydrolysis process, similar to previous theoretical studies (19, 20). We treat both the ATP and ADP-Pi actin as a single species denoted by ‘ADP-Pi.’ Thus, our model has three monomer states: ADP-Pi actin, ADP actin, and cofilin-bound ADP actin. We also consider the Pi-release to be random, i.e., ADP-Pi actin can convert to ADP actin with a rate *r*_1_ randomly over a filament length as reported before (21, 22). We modify a previously established model of ‘random hydrolysis’ (14, 19, 23–29) to incorporate the random severing events that crucially depend on the nucleotide states (ATP/ADP-Pi or ADP) of the monomers within the filament.

The severing proteins, ADF/cofilins, preferentially bind to the ADP actin monomers (30), with a rate *k*_*b*_ [*co f*]. It has been shown that cofilin binds to ADP monomers cooperatively (6, 8, 17, 31). Thus, following (6, 3 1), we assume that ADP actin converts to cofilin-bound ADP actin with a rate given by a hill function, 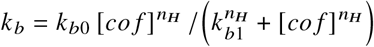, where *n*_*H*_ is the Hill coefficient which is a measure of cooperativity, *k*_*b*0_ and *k*_*b*1_ are positive real parameters. Additionally, recent papers have reported that cofilin binding enhances the Pi release of adjacent ADP-Pi monomers (6, 8, 17). We consider this process using a faster rate *r*_2_ (*r*_2_ ≫ *r*_1_) and assume a spatial range R (in units of monomers) around a cofilin-bound ADP monomer where the Pi-release may happen faster (similar to a model by C Suarez et al. (9)).

Experiments have suggested that cofilin binding induces mechanical stress discontinuities at the boundaries between ATP/ADP-Pi monomers and cofilin-bound ADP monomers (9, 32). Thus, we consider that the severing occurs with a rate *k*_*sev*_ at the interfaces of the undecorated monomers (i.e., cofilin unbound, comprising of ADP-pi and ADP actin) and cofilin-bound or decorated ADP monomers, generating two pieces: one still connected to the original barbed end and the other with a new barbed end (Fig 1C). Note that the filament length is measured from the original barbed end that is kept in focus (Fig 1C), while we assume that the fragment severed from the filament detaches and diffuses away from the focal plane (7). Moreover, since a large number of capping proteins exist in vivo (4, 17), the severed fragment could be capped rapidly and may not elongate.

Finally, if *l* represents the length of the filament (in units of monomers) as measured from the original barbed end, we can summarize all the kinetic processes by the following three reactions (Fig 1C):

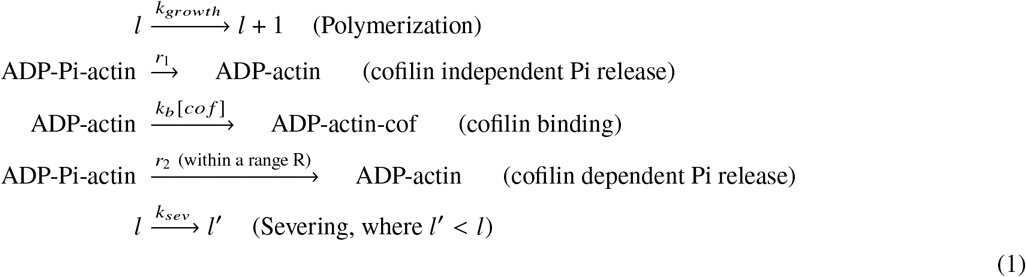

Here, note that severing is a zeroth order reaction and is proportional to the number of available severing sites in the filament (i.e., the number of interfaces between undecorated and cofilin-decorated ADP monomers). Thus, 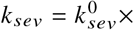 number of severing sites, where 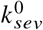is the bare severing rate.

### Simulation Method

We performed exact simulations of the processes described in Eq. (1) using the Gillespie (Kinetic Monte Carlo) algorithm (33). We use realistic parameter values obtained from in vitro experiments and previous studies to get the time evolution of the filament length (see Table 1 for parameter values). To calculate length distributions, we have simulated multiple stochastic trajectories of the length starting from an initial length of zero. To ensure a steady state, we sample from many trajectories at a large time so that the mean and variance of length become independent of time. Note that we express actin length in units of monomers for convenience; However, using the conversion factor 1 monomer = 0.0027 *μm*, we can express the length in real units (15, 20).

**Table 1:** Parameters used for simulations.

| Symbols | Values | Comments |
| --- | --- | --- |
| $r_0$ | $11.6 \mu\text{M}^{-1} \text{s}^{-1}$ | (36) |
| $r_1$ | $0.0019 \text{s}^{-1}$ | (37) |
| $r_2$ | $0.05 - 1.3 \text{s}^{-1}$ | Varied in this study |
| $R$ | $10 - 100$ | Varied in this study |
| $k_{b0}$ | $0.015 - 0.15 \text{s}^{-1}$ | Varied in this study |
| $k_{b1}$ | $1.0 \mu\text{M}$ | Assumed for this study. |
| $n_H$ | $3.5$ | Hill coefficient (6) |
| $k_{sev}^0$ | $0.00001 \text{s}^{-1}$ | (9) |

Complementary to the exact simulations, we have also developed a simpler two-state coarse-grained model in limiting conditions, where we used a mean-field mathematical framework based on previous studies (19, 20, 34, 35) to make mathematical predictions (see SI for details).

## RESULTS

### Model reproduces experimental observations of non-monotonic filament length with cofilin concentration

First, we consider the results from the severing assay by D. Pavlov et al. (7). In this study, the severing activity was inferred by measuring the percentage change in the mean length when actin filaments were treated with yeast cofilin. The authors reported a nonmonotonic behavior in length change, indicating the highest severing activity at an intermediate cofilin concentration and lower activity at both low and high cofilin concentrations (data shown in Fig 1A). In this experiment, the barbed ends of pre-formed filaments were attached to CapZ, a capping protein that inhibits polymerization at the barbed end. Also, data were taken under pre-steady state conditions. Thus, to qualitatively capture the transient phenomenon, we set the polymerization rate (*r*_0_) to zero and simulated the filament dynamics up to *t* = 2 minutes (before reaching the steady state), starting from an initial length (*l* (0)) of 4000 monomers (≈ 10 *μm*). We then calculate the mean length (⟨*l* (*t*)⟩) over an ensemble and define the percentage length change as 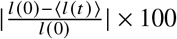. Remarkably, we obtain nonmonotonic curves for length change at different cofilin concentrations, as observed in experiments (compare Fig 2A and Fig 1A).

**Fig 2:**
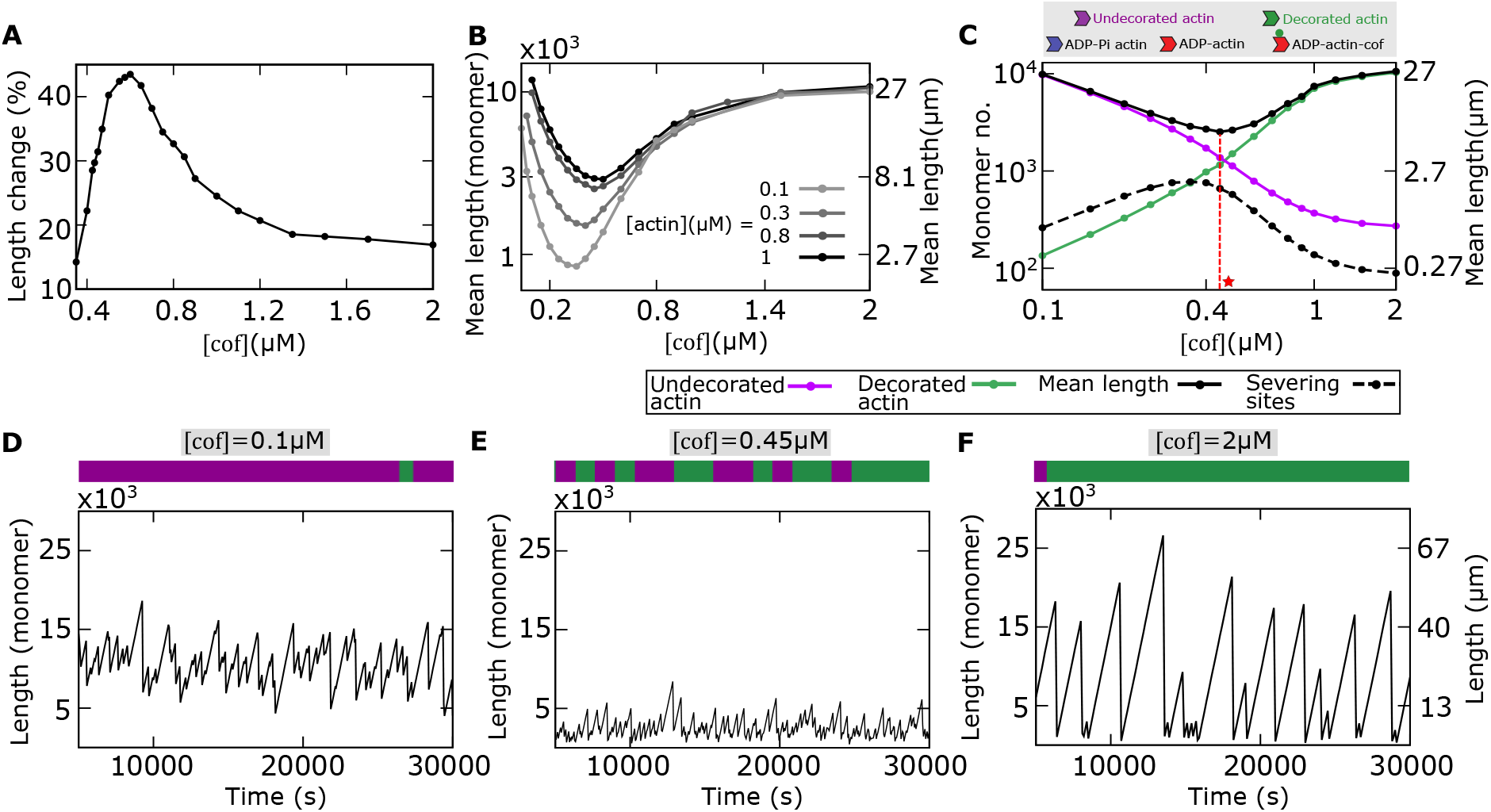
Model reproduces non-monotonic length changes seen in previous experiments. **(A)** Average length change of a filament in severing assay is non-monotonic with cofilin concentrations, as reported in (7). **(B)** The steady state mean length varies non-monotonically with the cofilin concentration, and the non-monotonicity increases as actin concentration is decreased. **(C)** Relationship between mean filament length and the average numbers of cofilin-undecorated actin (ADP-Pi actin and ADP-actin combinedly, in magenta), decorated-actin (cofilin-bound ADP actin, in green), and the average number of severing sites as functions of the cofilin concentration. At an intermediate cofilin concentration (shown as a red star), the maximum of severing sites and the minimum mean length coincide with the crossover between the average numbers of undecorated-actin and decorated-actin within the filament. **(D-F)** Length trajectories obtained from stochastic simulations, plotted at different cofilin concentrations. The microscopic arrangements of undecorated actin (magenta) and decorated actin (green) within a filament are shown above each trajectory. **(D)** At a low cofilin concentration, the filament is primarily composed of undecorated actins. The length trajectory is dominated by growth, with considerable fluctuations due to severing. **(E)** At an intermediate concentration, the heterogeneity in the arrangement of undecorated and decorated actin is more pronounced in the filament, with a lower mean length and smaller fluctuations. **(F)** At a high cofilin concentration, the filament is mainly composed of decorated-actins, exhibiting large length fluctuations due to sudden shrinkages, arising from more sporadic severing. Parameters: [*actin*] = 0.8*μM, r*_2_ = 0.19*s*^−1^, *k*_*b*0_ = 0.15*s*^−1^, *R* = 10. Other parameters: see Table 1.

Pavlov’s experiments were done at pre-steady state conditions, but with our theoretical model, we can also predict the mean length with cofilin concentrations at the steady state (Fig 2B). The convex nonmonotonic nature of the mean length at the steady state indicates that the severing activity is highest at an intermediate cofilin concentration, as observed in experimental studies (6, 8, 38). Thus, our model reproduces a nonmonotonic change in the mean length with the cofilin concentration. Since severing activity peaks at an intermediate cofilin concentration, we hypothesized that the internal arrangement of cofilin-bound and unbound monomers within a filament vary with cofilin concentrations. We study this in the next section.

### Non-monotonic behavior in length arises from the internal heterogeneity of monomer arrangements

To understand the origin of nonmonotonicity in the mean filament length, we now probe the arrangement of undecorated (i.e., ADP-Pi and ADP monomers combinedly) and decorated (cofilin-bound) monomers within a filament (Fig 2C). Using stochastic simulations, we calculate the mean numbers of decorated and undecorated monomers, the number of severing sites (the interfaces between decorated and undecorated actin monomers) and the resulting mean filament length as functions of the cofilin concentration. We find that the mean number of undecorated monomers monotonically decreases, while the mean number of decorated monomers increases with the cofilin concentration (Fig 2C). This is because the conversion of undecorated to decorated monomers increases with the cofilin concentration (Fig 2C). At an intermediate cofilin concentration of 0.4 *μM*, the mean numbers of undecorated and undecorated monomers become comparable on average. At this point, there is a higher number of interfaces between undecorated and decorated monomers, leading to a maximum for severing sites and a minima in filament length (Fig 2C). Interestingly, a maxima in severing activity (measured as *severing events μm time*) at an intermediate cofilin concentration was reported in a previous experiment in fission yeasts. (9). Our analysis sheds light on how that arises from the arrangement of monomers within a filament.

To further understand this behavior, we compare the actin length trajectories alongside the microscopic arrangements of undecorated and decorated monomers at low, intermediate, and high cofilin concentrations (Fig 2D-F). At low cofilin concentrations, the conversion of undecorated monomers to decorated is minimal, and hence, the filament is made of mostly undecorated monomers with fewer severing sites (see the configuration in Fig 2D, top). This leads to pronounced filament growth with occasional shrinkage events due to sporadic severing, as observed in the length trajectory (Fig 2D). Conversely, at much higher cofilin concentrations, the enhanced conversion rate from undecorated to decorated monomers causes the filament to be mainly composed of decorated monomers, again with fewer severing sites (Fig 2F, top). This also results in fewer shrinkage events similar to the case of low cofilin concentration (compare Fig 2D and Fig 2F). However, at intermediate cofilin concentrations, when the filament contains, on average, almost equal numbers of decorated and undecorated monomers, the number of possible severing sites also increases, leading to maximum heterogeneity in the monomeric arrangements (Fig 2E, top). This increased heterogeneity in monomer composition results in many instances of sharp shrinkages caused by multiple severing sites (Fig 2E), leading to a dynamic instability-like trajectory observed in microtubules (28, 39). Moreover, the configuration shown in Fig 2E (top) exhibits long domains/clusters of decorated monomers, that arise due to the site-specific spatial cooperativity in cofilin binding as reported before (9, 40–43).

### Actin length distribution and length fluctuation can be shaped by cofilin concentration

By examining the stochastic growth trajectories at the steady state, we observed that the frequency of elongation and shortening events (driven by severing) depend on cofilin concentrations. The frequency of these events is reduced at both lower and higher cofilin concentrations, but the frequency is enhanced at an intermediate cofilin concentration — this suggests that the filament length distribution and the fluctuation properties can qualitatively change with cofilin concentration. Next, we studied the steady-state length distributions associated with the trajectories at different cofilin concentrations. Our simulations show that the length distribution is bell-shaped at a lower cofilin concentration, but it gradually changes to skewed long-tailed distributions with increasing cofilin concentrations (Fig 3A). Notably, our assumption of hydrolysis-dependent severing at the interfaces of undecorated and decorated actin produces long-tailed distributions similar to experimental observations (7). This is in contrast with a previous theoretical study that mainly predicted bell-shaped length distributions (17), arising from hydrolysis-dependent severing.

**Fig 3:**
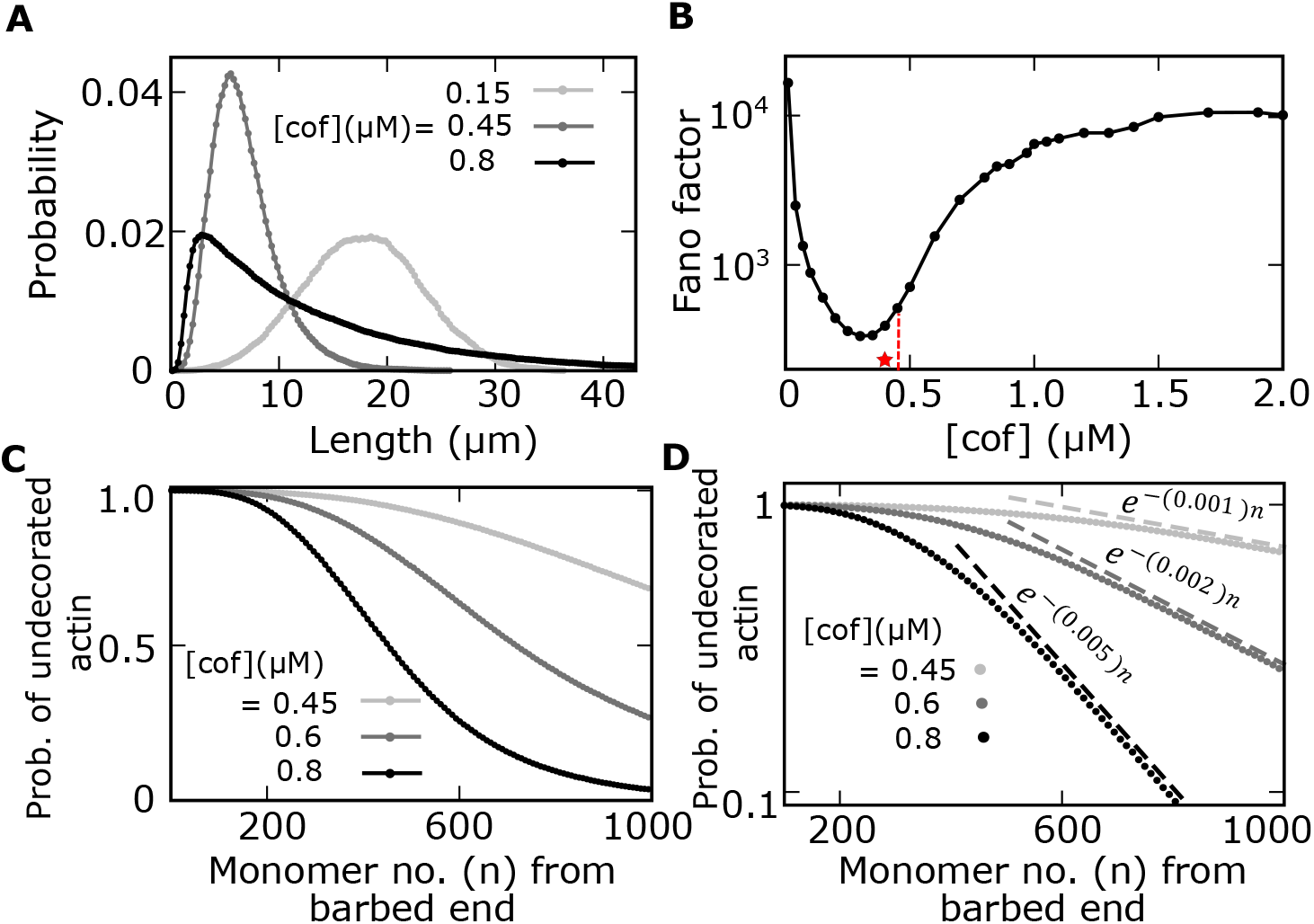
Distributions of Filament lengths and undecorated monomers within a filament. **(A)** Steady state filament length distributions vary from bell-shaped to skewed long-tailed shapes for different cofilin concentrations. **(B)** The minima of the Fano factor of the length distribution coincide with the intermediate cofilin concentration (denoted by a red star), corresponding to the minimum mean length in Fig 2C. **(C-D)** The probability distributions of cofilin-undecorated actin within a filament decrease with monomer numbers from the barbed end, shown at different cofilin concentrations. **(D)** Semi-log plot of the distributions of undecorated actin, showing that distributions have exponential tails (fitted exponential curves shown in dashed lines). Parameters: [*actin*] = 0.8*μM, r*_2_ = 0.19*s*^−1^, *k*_*b*0_ = 0.15*s*^−1^, *R* = 10. Other parameters: see Table 1.

To further quantify the length distributions, we calculate the Fano factor (ratio of variance to the mean) and find it varies non-monotonically with cofilin concentration (Fig 3B). Interestingly, the minima of the Fano factor is at an intermediate cofilin concentration close to the concentration where the mean length is minimum (denoted by stars in Fig 2C and 3B). This shows that noise associated with the filament distribution is correlated to the internal heterogeneity within the filament.

Moreover, we find that the probability of undecorated monomers (cofilin unbound monomers) within a filament rapidly decreases with the monomer number from the barbed end (Fig 3C). Intuitively, actin monomers near the barbed end are more likely to be in the ADP-Pi state than the monomers further away since ‘aged’ monomers are more prone to undergo hydrolysis and become ADP monomers on which cofilins can bind cooperatively. Thus, the probability of undecorated monomers should be a decreasing function with the monomer number. Significantly, we note that these probabilities have exponential tails that depend on cofilin concentrations (Fig 3D). In order to understand the origin of exponential tails, we next examine the arrangement of ADP-Pi, ADP, and cofilin-bound ADP monomers within a filament at different cofilin concentrations.

### Cofilin concentration can tune the arrangement of monomer states within a filament

In this section, we investigate how different model parameters affect the arrangement of monomer states within a filament as a function of the cofilin concentration. We begin by plotting the average numbers of individual ADP-Pi actin (blue), ADP-actin (red), decorated-actin (cofilin-bound ADP-monomers in green), and the undecorated-actin (ADP-Pi and ADP-actins combinedly, in magenta), as functions of the cofilin concentration, along with the corresponding mean length and the average number of the severing sites. With various parameter choices, we generally find the mean length exhibits a minimum, and the corresponding number of severing sites shows a maximum at an intermediate cofilin concentration where the average number of undecorated and decorated monomers cross each other (Fig 4A-D). For all chosen parameters, the average number of decorated (cofilin-bound) actin always increases, while both the numbers of ADP-Pi and ADP monomers decrease with increasing cofilin concentration (Fig 4A-D). Despite these qualitatively similar results across different parameter regimes, the relative abundance of the three monomeric states (ADP-Pi, ADP, and cofilin-bound ADP actins) can be varied by tuning specific parameters.

**Fig 4:**
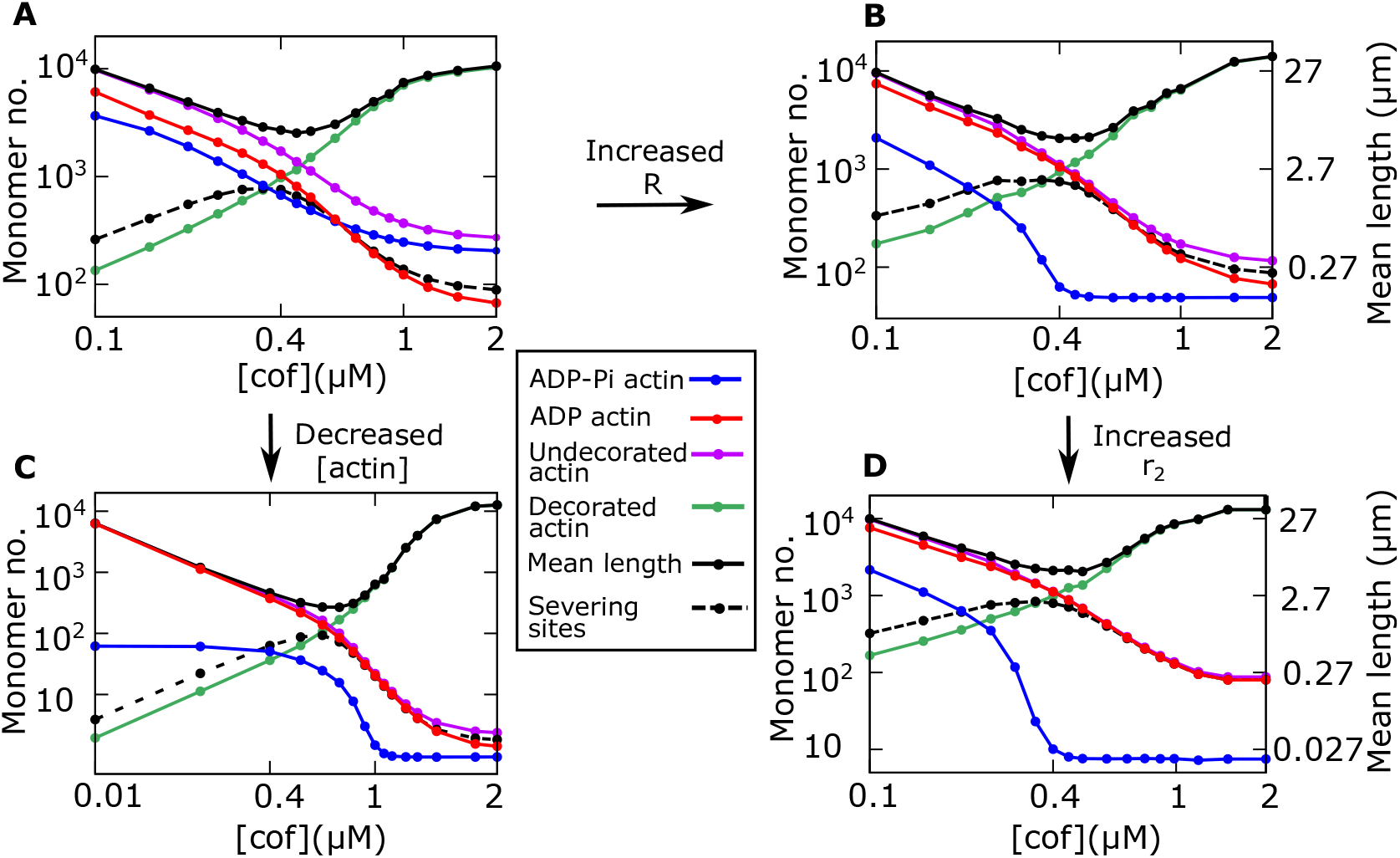
Arrangement of monomer states within a filament as a function of cofilin concentration. **(A-D)** Average number of individual ADP-Pi actin (blue), ADP-actin (red), decorated actin (cofilin-bound ADP-actin in green), undecorated actin (ADP-Pi actin and ADP-actin combinedly, in magenta), as functions of the cofilin concentration. The average number of severing sites (dashed line) and mean filament length (black) are also shown. By increasing range *R* from 10 (A) to 100 (B), lowering actin concentration from 0.8*μM* (A) to 0.01*μM* (C), and increasing *r*_2_ from 0.19*s*^−1^ (B) to 1.3*s*^−1^ (D), the average number of undecorated-actin becomes closer to the average number of ADP-Pi actin, indicating that the filament is mainly composed of two monomeric states, ADP-actin and cofilin bound ADP-actin. Other parameters: *k*_*b*0_ = 0.15*s*^−1^, and see Table 1.

Interestingly, we find that in the limits of a very large spatial range of cofilin-dependent Pi-release (*R*), at lower actin concentrations ([actin]), and for a very fast cofilin-dependent Pi-release rate (*r*_2_), the average number of undecorated actin becomes closer to the average number of ADP-Pi actin, essentially indicating that the filament is mainly composed of two monomeric states: ADP-cofilin-actin and ADP-actin (Fig 4B, 4C, and 4D). This result can be intuitively understood by noting that two processes mainly tune the relative abundance of ADP-Pi and ADP-actins: the influx of ADP-Pi actins via growth or polymerization and the conversion of ADP-Pi to ADP-actins via both cofilin-independent and cofilin-dependent Pi-release (see the Model in Fig 1C). Therefore, when the effective growth rate is much smaller than the Pi-release rates (i.e., *r*_0_ [*actin*] ≪ *r*_1_, *r*_2_), and when the spatial range representing the propagation of Pi-release allosterically from a decorated monomer is very high (*R* ≫ 1), the abundance of ADP-Pi monomers becomes very low across the filament as most of the ADP-Pi monomers quickly get converted into ADP-actins. In this case, the undecorated actins are mostly comprised of ADP-actins. This scenario motivated us to propose a coarse-grained two-state model that can mathematically show the origin of the exponential tails in the distribution of undecorated actins along the filament and also allows us to make analytical predictions about the actin cap size.

### A coarse-grained model can mathematically predict actin cap length variation

As described in the previous section, in some parameter regimes where the number of ADP-Pi monomers is substantially lower than the ADP-actins, we can reduce the original model into a simpler model consisting of only two monomer states – undecorated (cofilin-unbound) and decorated (cofilin-bound) actin, undergoing three processes, growth, cofilin-dependent switching from the undecorated to decorated actin, and severing at the interface of decorated and undecorated actins (Fig 5A).

**Fig 5:**
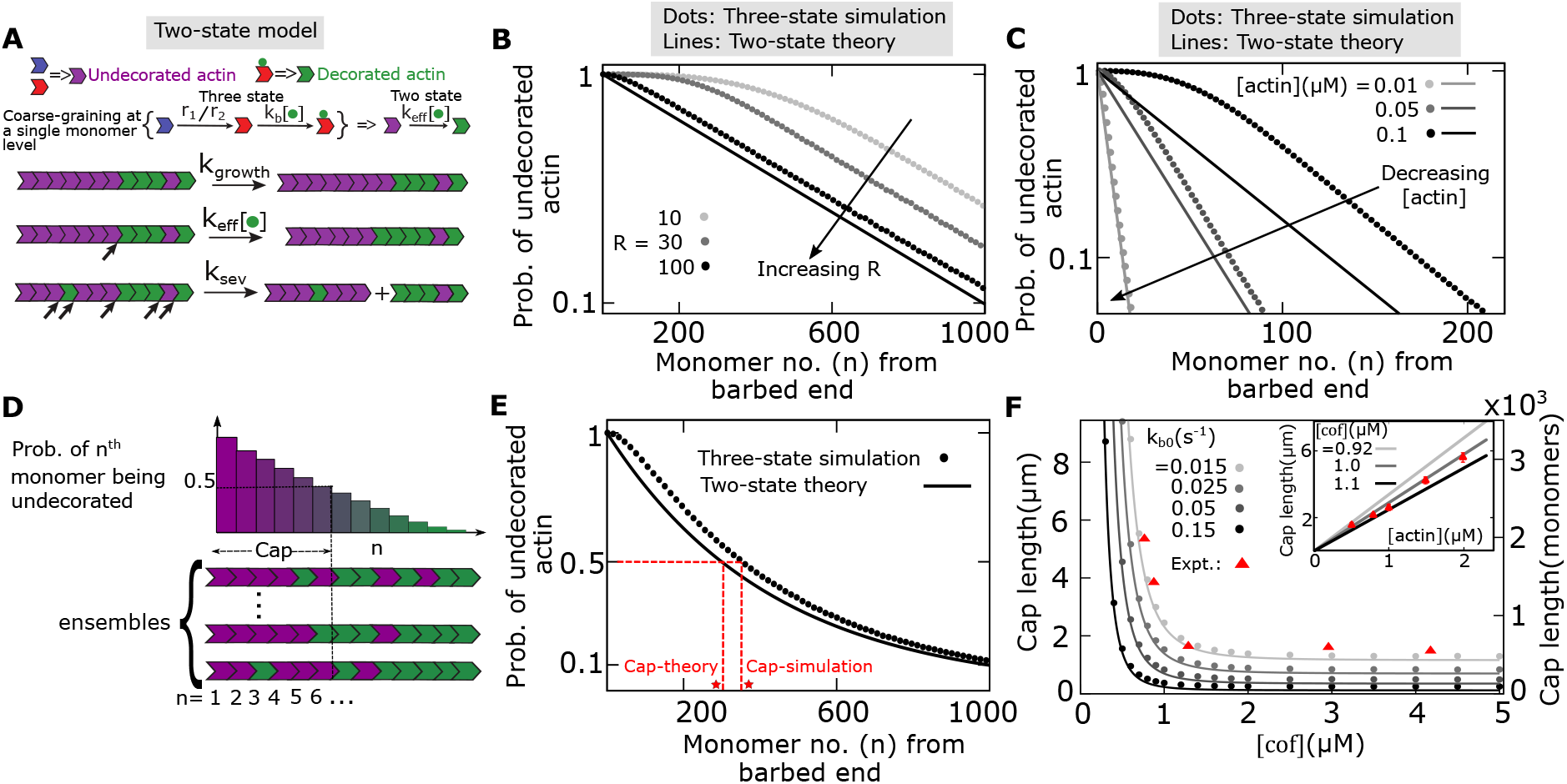
Coarse-grained model of hydrolysis-dependent severing reproduces actin cap size observations. **(A)** Two state model is made of two monomer types – cofilin-undecorated and cofilin-decorated actin, undergoing three processes, growth (with rate *k*_*growth*_), cofilin-dependent switching from undecorated to decorated (with rate *k*_*e f f*_), and severing (with rate *k*_*sev*_). **(B-C)** The probability of undecorated actin within a filament as a function of monomer number from the barbed end, shown for different values of (B) Pi release range *R* and (C) actin concentrations. Analytical predictions from the two-state model are shown in lines, and simulations from the three-state model are represented by dots. **(D)** Cap length is defined as the length from the barbed end, where the probability of an undecorated monomer becomes 0.5. **(E)** Capsize extracted from three state simulation reasonably matches with the value predicted by the two state model. Measures of the cap size from theory (cap-theory) and simulation (cap-simulation) are shown in red stars. **(F)** Variation of the cap size with the cofilin concentration for different values of the coarse-grained parameter *k*_*b*0_. Model reproduces previously reported results for the actin cap length variation as a function of cofilin and actin concentration (inset) (9). Lines represent analytical predictions from the two-state model, and dots represent simulations from the three-state model. Red points are published experimental data (9). Parameters: [*actin*] = 0.8*μM* (B, E-F), [*co f*] = 0.6*μM* (B-C, E), *r*_2_ = 0.19*s*^−1^ (B-C, E-F), *k*_*b*0_ = 0.15*s*^−1^ (B-C, E), 0.015*s*^−1^ (F (inset)), *R* = 10 (C), 100 (E-F), and for other parameters: see Table 1.

Note that, in the original model, the conversion from the ADP-Pi actin to decorated actin proceeds in two steps: conversion of ADP-Pi to ADP actin (via cofilin-independent or cofilin-dependent Pi-release), and then, the cooperative binding of cofilins to the ADP monomers. When the number of ADP-Pi actin is much smaller compared to the ADP actin within a filament, we can coarse-grain these two steps by a cofilin concentration-dependent effective switching from undecorated to decorated actin with a rate given by a Hill function, 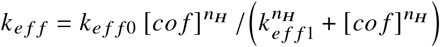, where *n*_*H*_ is the Hill coefficient. This assumption makes our model analytically tractable for mathematical predictions (see SI for details).

By solving the two-state model under the mean-field approximation, we derive an expression of *P*_*n*_, the probability of the *n*-th monomer from the barbed end being undecorated (see SI). We predict a geometric distribution given by *P*_*n*_ = *P*_1_*a*^*n*−1^, where the parameters *P*_1_ and *a* can be calculated in terms of other model parameters (see SI, Eq. S3). This provides a mathematical explanation of observing the exponential tails in the probability distribution of undecorated actins along the filament (Fig 3D). Interestingly, although derived from a simple two-state model, the predicted exponent of distribution tails matches quite well with the exact simulations of the actual three-state model (Fig 5B, 5C). In fact, at the limits of a high spatial range of cofilin-dependent Pi-release (high R), low actin concentration, and high cofilin-dependent Pi-release rate, the three-state simulation results for the probability of undecorated-actin match exactly with the mathematical prediction of the two-state model (Fig 5B, 5C, and S1).

To validate our model further, we compare our model predictions with the experimental measurements of the ATP-cap length of an actin filament, as measured by C. Suarez et al. (9). Typically, the actin cap refers to the continuous stretch of unhydrolyzed monomers (consists of ATP and ADP-Pi actins) from the barbed end, as defined in previous theoretical studies (16, 25, 44). However, resolving the fluorescence of a tagged ATP/ADP-Pi monomer at the microscopic level is experimentally challenging since small ADP islands inside a continuous stretch of ATP/ADP-Pi monomers may be undetectable. Hence, to match the cap length measured from the fluorescence intensity, we use the definition of Suarez et al. (9). In this study, the cap length was defined as the stretch of monomers (measured from the barbed end) within which the probability of finding undecorated monomers is greater than or equal to 50% (i.e., the value of *n* for which *P*_*n*_ = 0.5; see Fig 5D, 5E). Following this definition, we extract a capsize from the three-state simulation and compare it with the value predicted from the coarse-grained model (Fig. 5E). Remarkably, we find that the cap size from the three-state model simulation matches reasonably well with the prediction from the coarse-grained model when *k*_*e f f* 0_ = *k*_*b*0_ and *k*_*e f f* 1_ = *k*_*b*1_, suggesting that Pi releases rates (*r*_1_, *r*_2_) and the range of spatial cooperativity (*R*) are irrelevant in this parameter regime.

Next, we calculate capsize as a function of cofilin concentration (Fig. 5F). To fit the cap size with the experimental data (9), we vary the parameter (*k*_*b*0_) that tunes the coarse-grained conversion rate from undecorated to decorated actin (see Table 1). Remarkably, we find that our original three-state model as well as the simplified two-state model reproduce previously reported results for the actin cap length variation both as the functions of cofilin and actin concentrations (Fig 5F). We show that the average cap size monotonically and rapidly deceases with the cofilin concentration (Fig 5F), but it linearly increases with the actin concentration (Fig 5F, inset), as reported in experiments.

## CONCLUSION AND DISCUSSION

Severing proteins, such as cofilins, exhibit affinities to actin filaments depending on the nucleotide state of actin monomers. Here, we have developed a model to study the connection between nucleotide state-dependent severing and resulting actin length distributions. We reproduce previously observed experimental observations using a computational model of actin filament dynamics that combines stochastic polymerization, hydrolysis-dependent cooperative binding of cofilins, cofilin-induced enhancement of Pi release, and severing of actin monomers. In particular, our model captures the non-monotonicity in the mean filament length, the monotonic decrease of ATP/ADP-Pi cap length with the cofilin concentration, and a linear increase of the cap size with actin concentrations, as found in previous experiments (7, 9). Further, we also show that, in limiting conditions, our model reduces to a simpler coarse-grained model that can mathematically predict the cap size variation as a function of cofilin and actin concentrations.

Our study reveals a physical connection between the internal heterogeneity within a filament arising from random arrangements of undecorated (ADP-Pi and ADP) and cofilin-decorated monomers, and the emerging length fluctuations (Fig 2). At an intermediate cofilin concentration, higher heterogeneity in monomeric arrangement produces a maximum number of severing sites within a filament, leading to higher frequency of sharp shrinkages (similar to the dynamic instability of microtubules). Conversely, the microscopic variation in monomer arrangements is much lesser at lower and higher cofilin concentrations, producing a lower frequency of elongation and shrinkage events. Such predicted differences in trajectories can be tested by analyzing in vitro kymographs, which in turn can reflect the differences in the underlying heterogeneity of severing sites within a filament. Intriguingly, in line with our prediction, it was found that the measured number of severing events (in units of *severing events* / *μm* /*s*) maximizes at an intermediate cofilin concentration but stays minimal at both lower and higher cofilin concentrations (9). Also, at an intermediate cofilin concentration, higher length fluctuations suggest that the time to reach the steady state is faster than lower and higher cofilin concentrations, where severing-driven shrinkage events are rare. Thus, in vitro measurement of the time to reach the steady state could also be a means to test our theory.

Our model further shows that the maximum number of severing sites at an intermediate cofilin concentration signals a minimum in the mean filament length as a function of the cofilin concentration (Fig 2B). As mentioned before, this non-monotonic change in mean length aligns with the in vitro experimental observation. In addition, we predict that the non-monotonicity in length becomes sharper by decreasing actin concentration (Fig 2B).

Moreover, at a constant actin concentration ([*actin*] = 0.8*μM*), we predict different types of length distributions with variations of cofilin concentrations, ranging from peaked bell-shaped distributions to skewed long-tailed ones (Fig 3A). Interestingly, an in vitro study analyzing the interaction of cofilins with actin filaments reported skewed long-tailed length distributions (7). Our study reveals how one can tune the shape of distribution by changing the cofilin concentration. Contrastingly, another theoretical work (17) studying the interaction of cofilins with actin filaments predicted primarily bell-shaped length distributions. This result hinged on the assumption that severing occurs between two consecutive cofilin-bound ADP monomers. Instead, as shown in recent experiments (9), severing takes place at the interfaces between cofilin-undecorated and cofilin-decorated monomers. We have incorporated this observation into our model, which led to various length distribution shapes, not just bell-shaped ones.

We make another testable prediction that the length fluctuations quantified by the Fano factor of the distribution has a minimum close to the cofilin concentration where the number of severing sites is maximum or the mean length is minimum (Fig 3B). Of note, quantitative measurement of the noise in distributions has been used to understand the molecular processes in the studies on gene expression noise (45–48). However, this strategy has not been utilized to understand the regulation of filament length variation. Here, we propose and show that quantification of length fluctuations (captured by the Fano factor) can provide clues about the underlying mechanism of length control in actin filaments. Further, we also predict that the proportion of undecorated monomers along a filament will decay exponentially (Fig 3C, 3D). Our simple coarse-grained model also provides a mathematical formula of this decay exponent as the function of actin and cofilin concentrations, which may be tested using experimental assays developed by Suarez et al. (9).

In summary, our study shows how tuneable parameters such as the cofilin concentration can produce a variety of length distributions for actin filaments. This general understanding can be applicable to other types of severing proteins that act on cytoskeletal filaments. For instance, microtubules also display GTP-dependent severing (49–51). Our theoretical framework can be adapted for microtubules to draw crucial predictions about hydrolysis-dependent severing in microtubules. Our study also highlights how non-equilibrium processes can modulate structural heterogeneity at the monomer level within a filament to shape the length variation in actin structures.

## ACKNOWLEDGMENTS

This work was supported by the National Institute Of General Medical Sciences of the National Institutes of Health R35GM147556 and Rochester Institute of Technology start-up funds (SR and LM). DD and BB also acknowledge the financial support from the Science and Engineering Research Board (SERB), Government of India (Grant number: EEQ/2023/000551), and IISER Kolkata.

## Supplementary Information

### S1 CALCULATION OF *P*_*n*_

Here we have used a mean field mathematical framework following previous studies (1–4). We first define a variable *P*_*n*_ as the probability that the n-th monomer is undecorated, and hence (1 − *P*_*n*_) is the probability that the *n*-th monomer is decorated. We can then write down the time evolution equations (Master equations) of these state probabilities and solve them to obtain an analytical expression for *P*_*n*_. Thus, the Master equations for *P*_*n*_ would be, for *n* = 1,

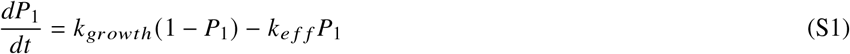

and, for *n* ≥ 2,

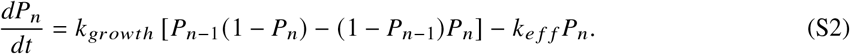

These equations are written by collecting the ‘gain’ and ‘loss’ terms for the probabilities. Also, note that adding an ADP-Pi monomer via polymerization shifts the site indices in the terms since the reference frame for counting the *n*-th site starts from the barbed end. For instance, in Eq. (S2), the addition of an ADP-pi monomer at the barbed end can positively contribute to the *n*-th site if the configuration was *P*_*n*−1_ (1 −*P*_*n*_). However, it negatively contributes to *P*_*n*_ if it was (1− *P*_*n*−1_)*P*_*n*_ before adding a monomer. Note that, *k*_*e f f*_ is the coarse-grained effective switching rate from an undecorated (ADP-pi or ADP) to a decorated (ADP-actin-cof) monomer. Additionally, this switching always changes the *n*-th site from *P*_*n*_ to (1 − *P*_*n*_).

In the steady state, the above equations (Eqs. S1 and S2) reduce to 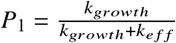 and 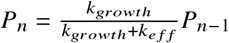. Therefore, we obtain

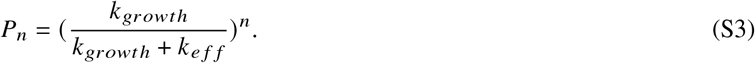

Note that *P*_*n*_ is a geometric distribution.

### S2 SUPPLEMENTAL FIGURES

**Figure S1:**
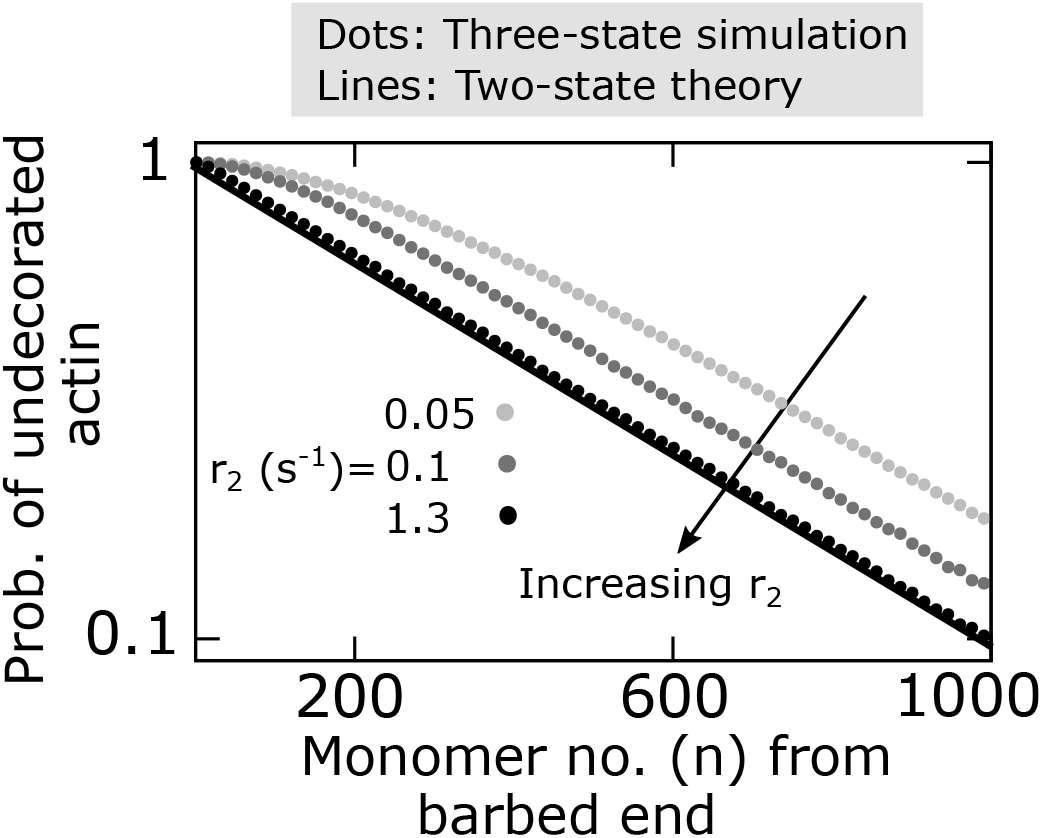
The probability of undecorated-actin within a filament as a function of monomer number from the barbed end, shown for different values of cofilin-dependent Pi release rate *r*_2_. Analytical curves from the two-state model are shown in lines, simulation from the three-state are represented by dots. Parameters: [*actin*] = 0.8*μM*, [*co f*] = 0.6*μM, k*_*b*0_ = 0.15*s*^−1^, *R* = 100. Other parameters: Table 1.

